# Bacterial strains in the human gut associate with host physiology

**DOI:** 10.1101/2025.11.30.691432

**Authors:** Saar Shoer, Anastasia Godneva, Adina Weinberger, Katherine S Pollard, Yitzhak Pilpel, Eran Segal

## Abstract

The human microbiota influences host physiology, yet much of its complexity lies beneath the species level. Here, we analyzed the intra-species genetic diversity of 936 gut bacteria across 24,997 individuals from three countries. Our findings show that highly abundant species exhibit greater strain stability, whereas low-abundance species display increased in-clonal mutations. Clonal strains are often mutually exclusive, while genetically variable strains tend to coexist. Strain turnover is associated with the presence of annotated chemotaxis and sporulation genes in reference genomes, whereas strain coexistence is associated with quorum sensing and secretion systems. Leveraging deep phenotypic data, we constructed an atlas detailing strain-level associations with diverse host physiological domains. For example, *Phocaeicola vulgatus* sub-types relate to host obesity, *Lachnospira eligens* to sleep, and *Parabacteroides distasonis* to iron hemostasis. This resource may guide personalized microbiome-based interventions to improve human health.

## Introduction

Bacteria inhabiting the human body harbor orders of magnitude more genes than the human genome, offering substantial genetic diversity and vast functional potential to influence host physiology^1^. Indeed, over a billion years of microbial-mammalian co-evolution has led to interdependency between the microbiota and its host. For example, gut bacteria, at the species level, have been associated with host beneficial outcomes such as the ability to digest certain foods, but also with a wide range of host diseases^2^.

Nonetheless, two strains of the same bacterial species can differ in their genetic makeup, resulting in pronounced phenotypic differences in both the bacteria and the host^3^. Recent studies have begun to demonstrate that even subtle intra-species genetic variations within human gut bacteria–including structural variations, presence or absence of accessory genes, and single nucleotide polymorphisms (SNPs)–are associated with host characteristics such as age, sex, and body mass index (BMI), leading to mechanistic insights^4–6^.

The bacterial subspecies population structure is shaped by the balance between clonal reproduction and homologous recombination^7–9^. Linkage disequilibrium (LD), the non-random association of alleles at different loci, provides a measure of this balance. Species with high LD typically exhibit low recombination rates, leading to the persistence of distinct, stable strain lineages. In contrast, low LD reflects frequent genetic exchange, which reduces associations between alleles and blurs strain lineage boundaries. Recurrent mutations can also depress LD. Thus, patterns of LD and SNP dissimilarity can reveal the mode and tempo of strain divergence.

Unlike the host (e.g., the human) genome, the genetic makeup of the gut microbiota is modifiable through strain losses and gains as well as via evolution of existing strains^10^. These genetic changes can be triggered by interventions such as diet, lifestyle, probiotics, and antibiotics. However, precise manipulation requires two key advances: a deeper understanding of the ecological and evolutionary forces shaping the microbiota at the strain level, across many species, and a catalog of bacterial genetic variations associated with host physiology^11^.

Here, we analyzed the intra-species genetic diversity of 936 gut bacteria across 24,997 individuals from Israel, the Netherlands, and the United States (US). Our strain analyses provide unique insights into bacterial population dynamics that are not captured by the species level. For example, SNP and gene divergence appear to be coupled in some species but decoupled in others. An individual can be identified by their gut microbiome after four years with >94% accuracy using strain typing, but in <30% of cases using species abundance. Leveraging deep phenotypic data, we constructed an atlas detailing strain-level bacterial associations with 14 host physiological systems. This resource may guide personalized microbiome-based interventions to improve human health.

## Results

### Abundant species exhibit greater strain stability, whereas low-abundance species display increased in-clonal mutations

We studied the intra-species genetic diversity of human gut bacteria by measuring SNP dissimilarity of 936 bacterial species across pairs of metagenomic samples (methods). We utilized 29,586 fecal samples from 24,997 human participants (43.59% male, mean age 50.71 years, mean BMI 26.31kg/m^2^) from Israel^12^, the Netherlands^13–16^, and the US^17^, including 4,589 longitudinal samples collected two and four years apart (Table1).

**Table1.** Baseline characteristics of human participants. Abbreviations: std - standard deviation, BMI - body mass index. *One individual contributed 86 samples over two years, plus three additional samples outside this time window.

| Dataset<br>(n or mean±std<br>[range]) | Human<br>Phenotype<br>Project<sup>12</sup> | Ullman | Lifelines-DMP<sup>13</sup> | Lifelines-DEEP<sup>1</sup><br>4–16 | DayTwo<sup>17</sup> |
| --- | --- | --- | --- | --- | --- |
| Country | Israel | Israel | Netherlands | Netherlands | United States |
| Participants at<br>baseline | 9,600 | 1 | 8,178 | 1,130 | 6,088 |
| 2-years follow-up | 3,220 | 86* | 0 | 0 | 0 |
| 4-years follow-up | 948 | 3* | 0 | 333 | 0 |
| Age [years] | 51.74±7.62<br>[40–70] | 43 | 48.43±14.79<br>[8–84] | 48.20±11.70<br>[18–80] | 52.63±12.82<br>[13–93] |
| Sex [%male] | 46.88% | 100% | 42.05% | 41.76% | 40.80% |
| BMI [kg/m<sup>2</sup>] | 26.05±4.15<br>[15.35–51.04] | 22.74 | 25.55±4.39<br>[13.10–63.70] | 25.4±4.08<br>[17.6–43.3] | 27.91±6.47<br>[12.93–68.10] |
| Bacterial species | 706 | 339 | 572 | 581 | 505 |

SNP dissimilarity within species (measured as Jaccard dissimilarity of alleles across whole-genome co-covered positions) only weakly correlates with SNP LD (90^th^ percentile of squared Pearson correlation between major allele frequencies of SNPs separated by 1,000 base pairs), gene variability (Heap’s law exponent fitted to pangenome growth curves, describing how the number of unique genes increases as more strains are introduced)^5^, species relative abundance within individuals (normalized mean coverage over the 50% most densely covered regions in reference genome)^17,18^, and prevalence across individuals (−0.33≤Spearman r≤0.33, Bonferroni-adjusted p-value, from here after denoted as q<0.05, methods, Figure1). This suggests SNP dissimilarity provides unique insights into bacterial population growth and dispersal beyond traditional metrics.

**Figure 1.**
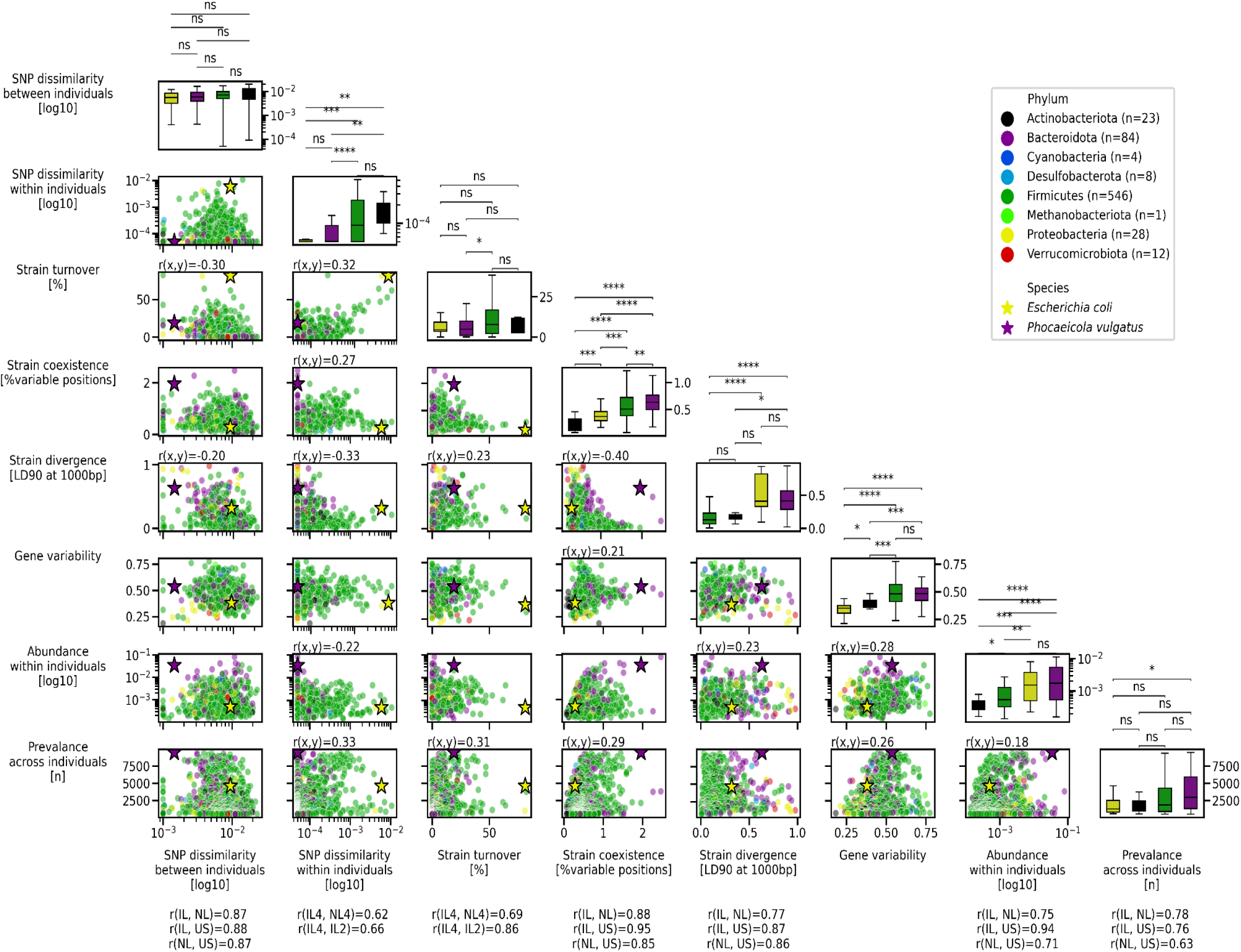
Baseline characteristics of bacterial species. Each row and column represents a different parameter measured in 706 human gut bacterial species of the Israeli dataset. From left to right and top to bottom, the parameters are: single nucleotide polymorphism (SNP) dissimilarity between different individuals at baseline; within the same individual after two years (note the different scales); percentage of individuals with strain turnover after two years; percentage of genomic positions with more than one allele (indicative of strain coexistence); 90^th^ percentile of SNP linkage disequilibrium (LD) at 1,000 base pairs (bp) (indicative of strain divergence); gene variability; relative abundance within individuals; and prevalence across individuals. Each dot in the scatter plots represents a species, colored by its phylum (legend, n indicates number of species). *Escherichia coli* and *Phocaeicola vulgatus* are highlighted with stars. Spearman correlations between the corresponding row and column parameters are shown above significant plots (q<0.05). Cross-country consistency among Israel (IL), the Netherlands (NL), and the United States (US) is indicated by the Spearman correlation below each parameter (q<0.05). “2” and “4” refer to the number of years between longitudinal samples. Box plots show the distribution of each parameter across the four major human gut bacterial phyla, ordered by median value. Boxes indicate quartiles; whiskers extend to 1.5 times the interquartile range. Outliers beyond this range are omitted. Mann–Whitney U tests annotations: not significant (ns) q>0.05, *q<0.05, **<0.01, ***<0.001, ****<0.0001.

All parameters show strong consistency across the three countries (0.62≤r≤0.95, q<0.05), indicating that human genetics and geographic dispersal within Western populations have limited effect on bacterial population dynamics. This conclusion is further supported by prior studies showing strains from European and North American samples tend to cluster together, and that the non-synonymous to synonymous polymorphism rates are more variable between gut microbial species than across human hosts^19,20^.

Using the baseline samples, the first time point samples of each country, we defined the typical SNP dissimilarity between individuals per species. Longitudinal analysis showed minimal SNP changes within individuals over years. In order to study strain stability, we defined strain turnover as cases where intra-person SNP dissimilarity exceeds the lower 5^th^ percentile of the inter-person distribution. We found that strain turnover is more frequent in species with high LD (r=0.23, q<0.001) and high prevalence (r=0.31, q<1e-7), independently, as these two parameters are not associated with each other (q>0.05). High LD indicates pronounced strain divergence. Strain replacement may explain the sudden (in evolutionary scale) and vast SNP shifts within individuals. High prevalence across humans may suggest the host community is the source of the new strains.

We next investigated whether the bacterial parameters differ across the four major human gut phyla–Firmicutes, Bacteriodota, Protobacteria, and Actinobacteria (n≥20 species, Figure1). While Bacteroidota and Proteobacteria exhibit the highest abundance and LD, they show the least SNP dissimilarity within individuals over time (Mann–Whitney U test q<0.05). This suggests that ecological adaptations such as niche specialization, competitive exclusion, or reduced selective pressure may underlie their genomic stability at the SNP level. At the gene level, Bacteriodota display the highest variability, whereas Proteobacteria display the lowest. This could indicate that SNP and gene divergence are coupled in some species, but decoupled in others, in line with a recent study showing the level of SNP and gene coupling varies across species^21^. In contrast, Firmicutes and Actinobacteria exhibit the lowest abundance and LD but the highest SNP dissimilarity within individuals over time, yet below the strain turnover threshold. This pattern may reflect either heightened mutation rates in response to increased selective pressures or reduced efficiency of purifying selection resulting from smaller effective population sizes. To rule out the possibility that the observed increase in in-clonal mutations among low-coverage species is an artifact of sequencing, we examined the relationship between species abundance at baseline and SNP dissimilarity after two years in each species across individuals with varying abundances. This correlation is minimal (r=0.019±0.17 for 429 species with q<0.05), supporting the conclusion that the increased in-clonal mutations observed in low-abundance species is a genuine long lasting bacterial trait.

Lastly, to identify mechanisms underlying variation in these bacterial parameters, we assessed whether the presence of annotated genes in reference genomes is associated with these parameters across species. We identified 1,227 associated genes (t test q<0.05), 257 of which are not explained by species taxonomic classification into the four major phyla (Chi-square test q<0.05, methods, FigureS1, Extended Data1). Most genes associate with only one parameter, indicating distinct selective pressures acting on different bacterial traits. Of the 36 genes whose presence relates to strain turnover after four years, 24 (67%), including 23 of the top 25, are linked to chemotaxis and the flagellum. The presence of ten genes containing CheY-like response regulator receiver domains is linked to species prevalence, five of which are directly linked to chemotaxis, while the remaining five, to the best of our knowledge, are not. Only one gene, *phoP*, a transcriptional regulator involved in virulence, environmental tolerance, and stress response^22^, is shared with the strain turnover associated set. These results suggest a role for chemotaxis–the directed movement toward favorable chemical stimuli via modulation of flagellar rotation–in strain turnover in the human gut.

### Clonal strains are often mutually exclusive, while genetically variable strains tend to coexist

Beyond how strains replace each other over time, a fundamental question is how they share the same ecological niche, evident from the high proportion of genomic positions harboring more than one allele in the same host in some species, termed strain coexistence (Figure1). Strain coexistence varies across phyla, with Bacteroidota exhibiting the highest levels, followed by Firmicutes and Proteobacteria, while Actinobacteria show the lowest (q<0.05). Similar to strain turnover, coexistence is more frequent in species with high prevalence across individuals (r=0.29, q<1e-12). This potentially reflects a greater likelihood for new strains to be introduced to the gut from the host community. In contrast to strain turnover, coexistence is less frequent in species with high LD (r=-0.40, q<1e-16), indicating clonal strains (i.e., high LD) are often mutually exclusive, while genetically variable strains (i.e., low LD) tend to coexist.

To identify mechanisms underlying strain coexistence, we utilized the functional analysis mentioned before. We find 99 genes associated with strain coexistence across species (q<0.05), of which 32 also associate with species prevalence and 23 with LD, supporting an underlying relationship among these parameters (FigureS1, Extended Data1). The presence of five genes is jointly associated with all three parameters–high strain coexistence, high prevalence, and low strain divergence–highlighting potential mechanisms enabling stable multi-strain colonization. These include: *luxS*, a secreted quorum sensing molecule that modulates both cell density signaling and metabolic communication; *ltaE*, an enzyme involved in amino acid catabolism, possibly facilitating metabolic cross-feeding; *cheB*, a chemotaxis regulator, potentially enabling niche partitioning and sporulation under stress; *rpoS*, RNA polymerase sigma factor enhancing survival under stress, including via sporulation; and *yafQ*, a toxin component of a type II toxin-antitoxin system that induces bacterial stasis.

Additional notable genes associated with strain coexistence, but not necessarily with the other parameters, relate to secretion systems types I–IV and horizontal gene transfer, suggesting that inter-bacterial communication and genetic exchange may facilitate or be required for stable multi-strain colonization. Further supporting the genetic exchange signal is the increased gene variability in species with high strain coexistence (r=0.21, q<1e-3). Sugar and vitamin metabolism and transport as well as the membrane are also frequent annotations in strain coexistence associated genes. Together, these findings suggest coexisting strains occupy complementary metabolic niches, potentially enabling stable multi-strain colonization through resource partitioning. This extends isolate-based estimates of gut strain richness^23^ to uncultured species in vivo and nominates specific mechanisms that may underlie stable multi-strain colonization.

### Hosts are stable over time with subtle strain acquisition from the environment

We next examined the strain stability of gut bacteria from the host’s perspective. We defined strain persistence within individuals and strain sharing across individuals as cases where the SNP dissimilarity is below the lower 5^th^ percentile of the inter-person distribution.

We found that unrelated individuals share a strain in fewer than 4.00% (median) of shared species, with no significant differences across countries, city of residence, nor participant characteristics such as year of birth, age, or sex (t test q>0.05, Figure2A). In the Israeli cohort, family members with shared genetic background–siblings and parent–child pairs–shared 6.45% and 10.00% of strains, respectively. In contrast, genetically unrelated but cohabiting partners shared 25.00% of strains (q<1e-12). As participants were over 40 years old, they are unlikely to live with their siblings or parents. This suggests that environmental exposure has a stronger influence than host genetics on shaping the microbiota at the strain level, consistent with previous findings at the species level^24^.

**Figure 2.**
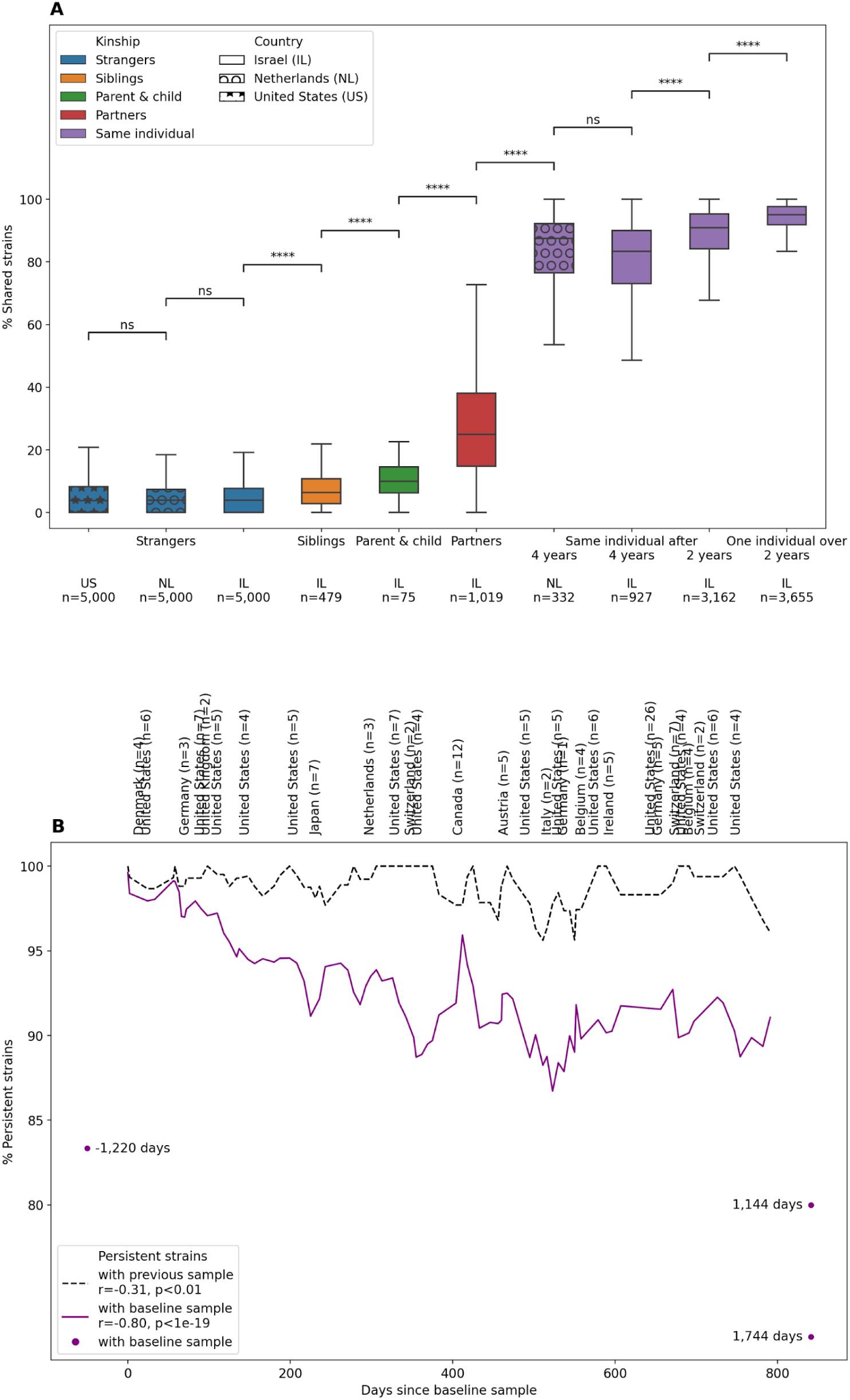
Hosts are stable over time with subtle strain acquisition from the environment. **A.** Percent of shared strains (y-axis) between individuals. Strangers (blue) are shown across three countries–Israel (IL, plain background), the Netherlands (NL, marked with circles), and the United States (US, marked with stars). Family members include siblings (orange), parent–child pairs (green), and cohabiting partners (red). Within-individual comparisons (purple) are shown after four and two years, with the rightmost box representing one individual’s comparison of 86 samples. Boxes indicate quartiles; whiskers extend to 1.5 times the interquartile range. t tests q-value annotations: not significant (ns) q>0.05, *q<0.05, **<0.01, ***<0.001, ****<0.0001. **B.** Longitudinal data from a single individual across time (x-axis), showing a three-point rolling mean of the percent of persistent strains (y-axis). Comparison to the previous sample is shown as a dashed black line; comparison to the baseline sample is shown as a solid purple line. Three additional samples taken outside the two-year window are plotted as purple dots with their actual time relative to the baseline indicated. Pearson correlations between strain persistent and days since the baseline sample are shown in the legend. International travels are marked above the plot, with trip durations (in days) indicated in parentheses.

To estimate how strain sharing decays over time after cohabitation ends, we used the older sibling’s age as a proxy for time since cohabitation and found a weak negative correlation with strain sharing (Spearman r=−0.12, p<0.01). Although weak, this correlation did not replicate when matching each older sibling with a stranger of the same age as the younger sibling (p>0.05). All tested species were more likely to be shared by cohabiting partners than by siblings (parent–child pairs were excluded from this analysis due to small sample size), but the two do correlate (r=0.33, p<0.01). *Faecalibacterium prausnitzii F*, *Prevotella copri B*, and *Sutterella wadsworthensis* showed the most pronounced differences, with strains shared in >60% of partners but <16% of siblings (FigureS2).

To identify mechanisms underlying environment-based strain sharing, we utilized the functional analysis mentioned before. We find 31 genes associated with the fraction of shared strains among partners across species (t test false discovery rate FDR-corrected p<0.05, FigureS1, Extended Data1). These include nine sporulation-related genes and five other stress-related genes. Notably, ten of these 14 genes are negatively associated with strain sharing.

Lastly, we used the longitudinal data to assess strain-level stability within individuals over time. In both the Dutch and Israeli cohorts, >83.33% of strains persisted in the same individual after four years, with no significant differences between countries, age, or sex (p>0.05, Figure2A, no longitudinal data was available for the US cohort). In the Israeli cohort, where two-year data was also available, strain persistence was higher at two years (90.91%) than at four years. In all cases, individuals with greater species richness exhibited higher stability (0.33≤Pearson r≤0.45, p<1e−17, FigureS3), as we showed in the past on a shorter time scale^25^. These results indicate that gut bacterial strains are largely stable over time, with minor gradual changes, in agreement with prior findings on smaller cohorts and smaller sets of species^16,19,26^.

This conclusion is further supported by high-resolution longitudinal data from one individual who contributed 86 samples over two years (mean interval 9.24±6.72 days, range 1–49), plus three additional samples outside this time window. From hereafter we call this unique cohort Ullman^27^. In this individual, the proportion of persistent strains is strongly tied with time from baseline sample (r=−0.80, p<1e−19), and only weakly tied when comparing each sample to the previous one (r=−0.31, p<0.01, Figure2B). The number of comparable species explained 22.61% of the variance in strain persistence relative to baseline (r^2^, p<1e−5), but only 6.37% relative to the previous sample (p<0.05). This suggests that reduced persistence at the species level over time is partially manifested at the strain level measurement.

More specifically, 98.85±1.75% (range 92.31–100.00%) of strains persisted between consecutive samples. From the baseline, 87.69% of strains persisted after two years, 80.00% after three, and 72.22% after five. On trend, 83.33% of the same strains were detected in a sample taken three years before the baseline, and 64.29% were still present eight years later. International travels had no significant effect on strain persistence (t test p>0.05). These findings further support the long-term stability of the human gut microbiota at the strain level.

### Strain typing is more consistent than species composition

We next compared dissimilarities at the strain level–both between individuals and within individuals over time–to dissimilarities measured at the species level. To do so, we converted the percent of shared strains into a strain existence dissimilarity (1−%shared strains/100) and computed species level existence and abundance dissimilarities using the Jaccard and Bray–Curtis methods, respectively (methods). Among the three approaches, the strain level showed the greatest variability across types of human kinships (FigureS4A).

Across familial kinships, strain-level measurements detected differences that species-level metrics often failed to capture (t test q>0.05). In contrast, across countries, in unrelated individuals and the same individual after four years, strain level measurements did not detect differences, while species level measurements did (q<0.05), potentially reflecting true biological variations across populations or technical artifacts caused by differences in sequencing protocols. In Ullman’s longitudinal samples, strain-level information showed minimal noise and gradual declining patterns over time, whereas species level measures often reported similar dissimilarity values between samples collected days and months apart, indicating reduced temporal sensitivity (FigureS4B).

To assess the accuracy of each method in identifying individuals over time, we compared the small sets of follow-up samples to the big sets of baseline samples and determined whether each follow-up sample’s closest match was the baseline sample of the same individual. Using strain existence we correctly identified 98.07% of Israelis after two years and 94.93% after four years. In comparison, species existence and abundance correctly identified only 71.74% and 29.23% of people after four years, respectively. Similar trends were observed in the Dutch cohort. The popular methods StrainPhlAn^28^ and MetaPhlAn 4.0^29^ based on marker genes showed 26.21% lower accuracy than our whole genome approach when using strain existence, and 4.53% and 2.58% lower accuracy when using species existence and abundance, respectively. StrainPhlAn can also estimate strain abundance; however, this metric performed considerably worse than strain existence (28.05% vs. 68.72%, TableS1). This shows strain typing by whole genome comparison conserves substantially more information than marker genes, albeit computationally more expensive.

The poor performance of species abundance potentially reflects true biological variation, as this feature was shown to fluctuate even within a day in certain species^30,31^. This fluctuation may also explain why species and strain existence, despite their lower resolutions, consistently outperformed species and strain abundance measurements, respectively. Overall, whole genome strain typing appears to be more consistent and informative than marker gene, compositional, and species-level approaches, which are more sensitive to collection timing and sequencing platforms. These findings further support earlier indications of this phenomena^19^ even in bigger sample sets, improved reference genomes, and more advanced measurement metrics.

### Different bacterial species exhibit different strain dynamics

We utilized Ullman’s high-resolution longitudinal data to investigate selected species strain-level dynamics in detail over a two-year period (Figure3 and S4). We focused on *Escherichia coli*, a model organism Proteobacteria species of moderate prevalence in humans, that typically appears in low abundance in our gut and exhibits exceptionally high levels of strain turnover across all longitudinal cohorts. To contrast this pattern, we chose to focus on *Phocaeicola vulgatus* (formerly *Bacteroides vulgatus*), a Bacteroidota species of high prevalence and abundance but only moderate levels of strain turnover in the longitudinal cohorts. Despite these differences, both species show pronounced strain divergence (Figure1 and S5). In Ullman, *E. coli* was detected in 27 samples, with abundance roughly one order of magnitude lower–and SNP dissimilarity approximately one order of magnitude higher–than *P. vulgatus*, which was detected in 85 samples. This aligns with the population level trends observed in these species across individuals, as well as the phylum-level results showing low-abundance species, such as *E. coli*, display increased in-clonal mutations.

**Figure 3.**
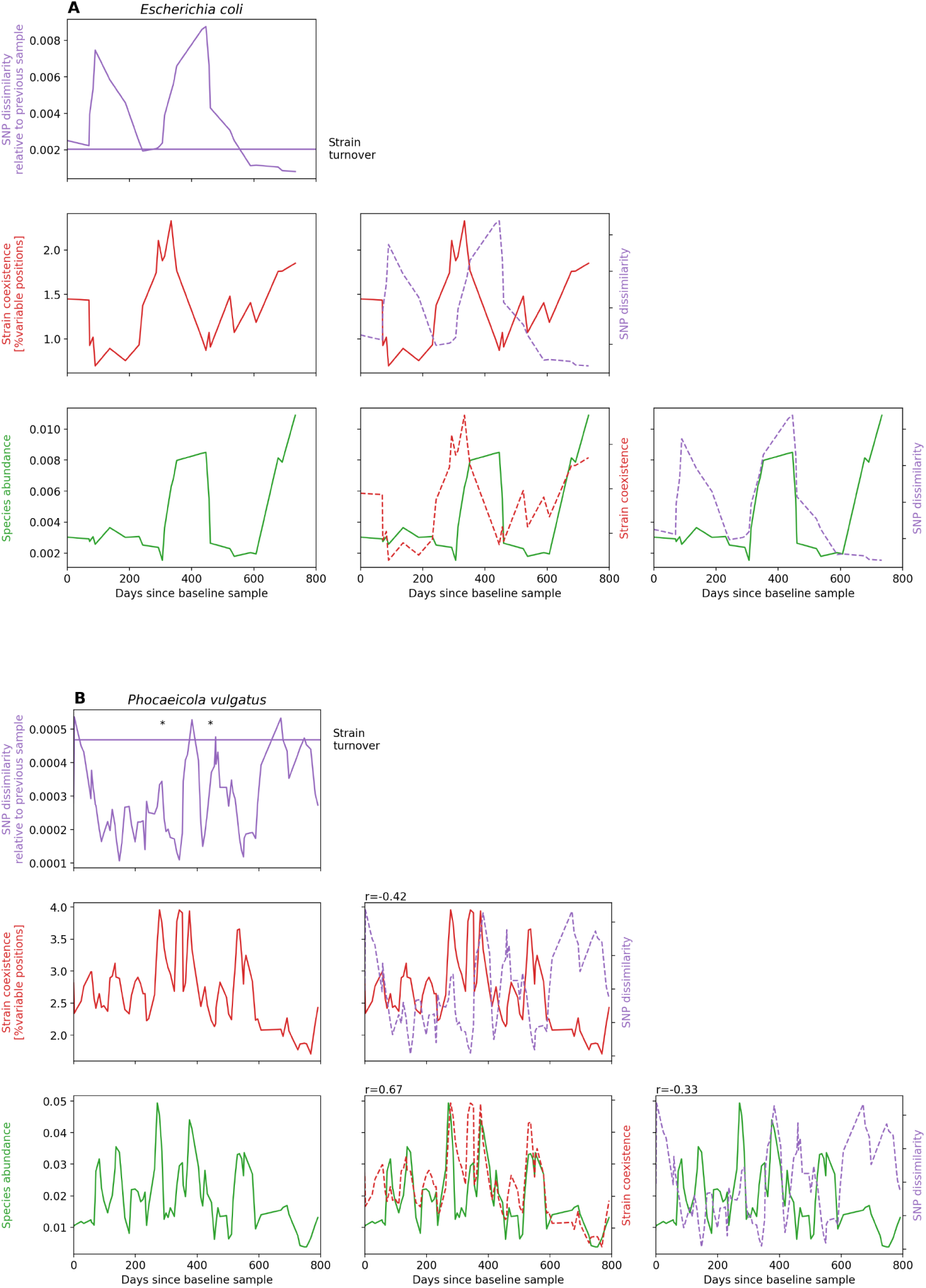
Different bacterial species exhibit different strain dynamics. Longitudinal profiles of *Escherichia coli* (**A**, n=27 samples) and *Phocaeicola vulgatus* (**B**, n=85 samples) from a single individual across time (x-axis). Each row corresponds to a different parameter (y-axis): single nucleotide polymorphism (SNP) dissimilarity relative to previous sample (purple), percentage of genomic positions with more than one allele (indicative of strain coexistence, red), and species relative abundance (green). Assembled genomes time points are indicated with asterisks. Overlaid plots show two parameters with dual y-axes (left and right). Spearman correlations between the two parameters are shown above significant plots (q<0.05).

Both species showed evidence of a rapid strain replacement event around day 400, preceded by an increase in variable positions, suggesting temporary multi-strain colonization before one strain per species became dominant. Interestingly, *E. coli* abundance peaked after the strain replacement, whereas *P. vulgatus* abundance peaked during the coexistence phase. We were able to assemble *P. vulgatus* genomes before and after the strain replacement and identified 1,925 genes shared between the two strains, with approximately 1,800 genes unique to each. Among 30 ribosomal protein genes we previously associated with younger hosts^5^, Ullman lost nine and gained two genes following the strain replacement event, consistent with host aging.

### Bacterial subspecies associate with host age, sex, and BMI

We checked whether different bacterial strains are associated with host characteristics. To this end, we defined subspecies types by clustering baseline samples of each species in each country based on SNP dissimilarity. Subspecies were delineated as groups of strains with high intra-group similarity and clear divergence from other groups (methods). SNP dissimilarity quantifies genetic distance between strains, whereas LD evaluates the balance between within-group cohesion and between-group separation. Accordingly, low LD yields more diffuse groups with shallower boundaries, whereas high LD favors tight, well-separated clonal clusters. Because these measurements probe different properties, they can vary independently. We found that different species exhibit distinct subspecies population structures described by these two parameters, and these structures were fairly consistent across the three countries (Figure4). For example, *Gemmiger qucibialis* has low LD and relatively uniform dissimilarity across participants, indicating minimal strain divergence. *Barnesiella intestinihominis* shows moderate LD and forms small, loosely defined lineages, with participants within each lineage highly similar to each other but different from participants in most other lineages. In contrast, *Parabacteroides distasonis* displays high LD and well-separated lineages, indicating high strain divergence.

**Figure 4.**
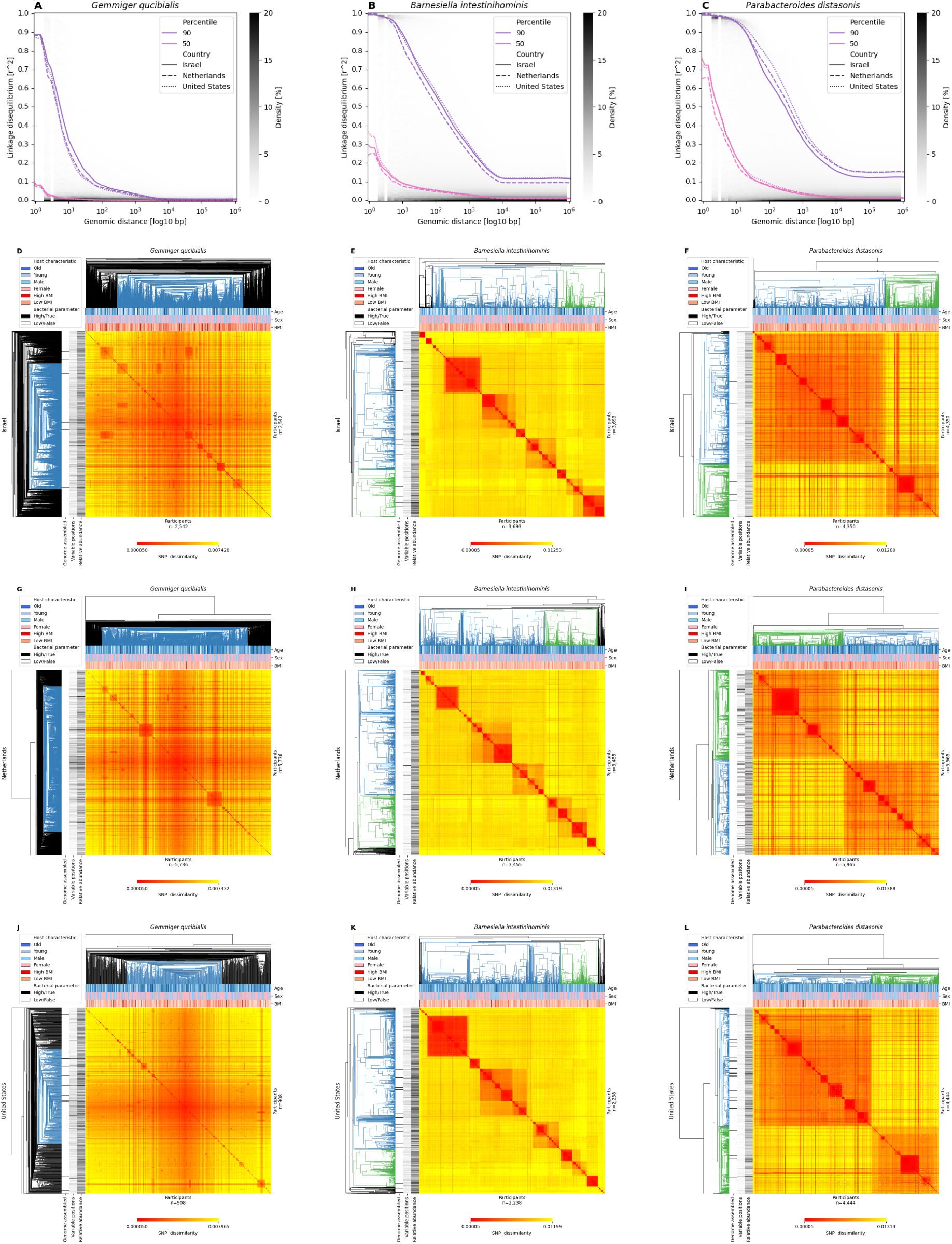
Different bacteria exhibit distinct subspecies population structures. **A-C.** 90^th^ (purple) and 50^th^ (pink) percentiles of linkage disequilibrium (LD, y-axis, calculated as Pearson correlation squared between single nucleotide polymorphisms - SNPs) as a function of genomic distance in base pairs (bp, x-axis) for *Gemmiger qucibialis* (A), *Barnesiella intestinihominis* (B), and *Parabacteroides distasonis* (C), in the Israeli (solid line), Dutch (dashed line), and American (dotted line) datasets. Grey background indicates data density. **D-L.** Cluster maps of SNP dissimilarity between participants at baseline for each species (columns) in the three countries (rows). Red-Yellow color scales indicate dissimilarity (bottom colorbars, note the different scales across species). Dendrogram colors denote sub-types. Horizontal colorbars above each heatmap indicate host characteristics; vertical colorbars to the left indicate bacterial parameters (legend).

Next, we tested for associations between bacterial sub-type presence and host age, sex, and BMI. Among 140 species tested in at least one country, 26 show association with age, ten with sex, and seven with BMI at the intra-species level, while controlling for the other two host variables (linear models q<0.05, methods). All but two associations remain significant after adjusting for species abundance, indicating the sub-type level adds information beyond species quantity. Notably, *P. vulgatus* is associated with all three host characteristics in all three countries, except for age in the US, which did not pass multiple hypothesis correction (only 31 species were tested in all countries, FigureS6, Extended Data2). In the Israeli cohort, participants carrying the *P. vulgatus* “blue” sub-type are, on average, 0.88 years younger, 12.59% more likely to be male, and have a 0.64 kg/m^2^ higher BMI than those carrying the “green” sub-type. Similar trends were observed in the other countries. Interestingly, a third group carries mixtures of both sub-types, indicated by low SNP dissimilarities from both the “blue” and “green” groups, increased variable positions relative to the other groups, and failure to assemble genomes, as strain mixture interferes with genome assembly procedure (Figure5).

**Figure 5.**
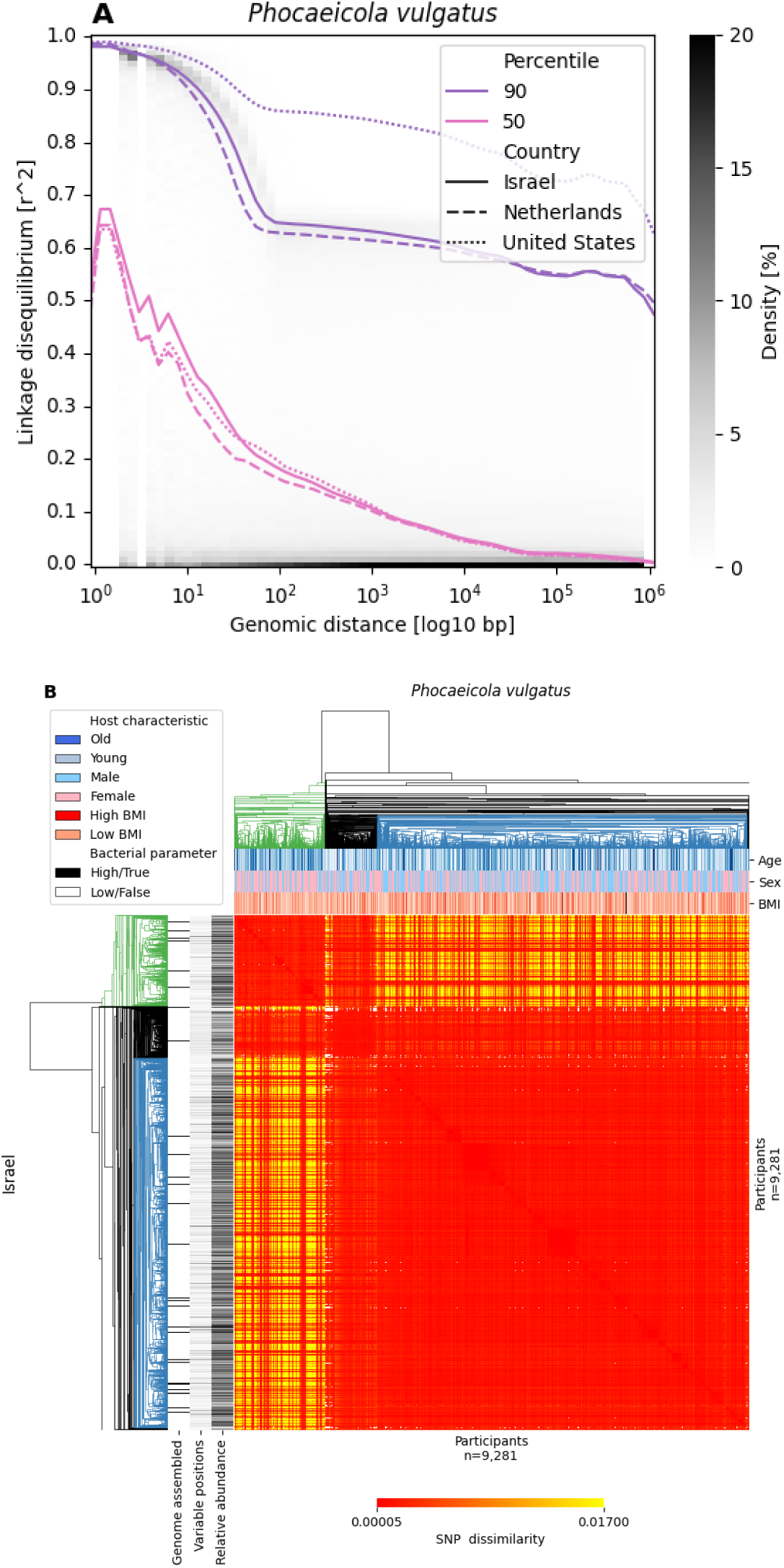
*Phocaeicola vulgatus* subspecies associate with host age, sex, and BMI. **A.** 90^th^ (purple) and 50^th^ (pink) percentiles of linkage disequilibrium (LD, y-axis, calculated as Pearson correlation squared between single nucleotide polymorphisms - SNPs) as a function of genomic distance in base pairs (bp, x-axis), in the Israeli (solid line), Dutch (dashed line), and American (dotted line) datasets. Grey background indicates data density. **B.** Cluster map of SNP dissimilarity between Israeli participants at baseline. Red-Yellow color scale indicates dissimilarity (bottom colorbar). Dendrogram colors denote sub-types. Horizontal colorbars above each heatmap indicate host characteristics; vertical colorbars to the left indicate bacterial parameters (legend).

Our previous study using gene presence/absence information from assembled genomes identified similar *P. vulgatus* “blue” and “green” sub-types and associations with host age and sex^5^ (1.67 years and 8.19% sex difference, BMI was not available), suggesting SNP and gene level divergence in this species are closely related. *Akkermansia muciniphila* also showed concordant sub-types at the SNP and gene levels and associations with host age and sex at the gene level. However, at the SNP level, only the age association replicated and only in the Israeli cohort (1.44 years difference; in the American cohort it was significant before the multiple hypothesis correction), implying that in this species SNP and gene divergence–particularly with respect to sex–may be less coupled (gene-level differences were 3.32 years and 7.73% sex). These observations align with the earlier phylum-level results that showed some phyla are more prone to SNP and gene coupling, while in others it is more often decoupled.

Other notable species include *Alistipes putredinis* and *Prevotella copri B.* Both are associated with host age across countries, although the association of *P. copri B* did not pass multiple hypothesis correction. Similarly, an unclassified Hominicoprocola species is associated with host’s sex and *P. distasonis* with BMI in all three countries but they are not significant after multiple hypothesis corrections. *E. coli* sub-types show no association with host characteristics. Low species prevalence across individuals reduces sample size and statistical power which may explain the lack of results in some species.

We performed similar association tests for two bacterial parameters: variable positions (indicative of strain coexistence) and species relative abundance within individuals. Sub-type-level differences were detected in at least one country for 55 and 54 species, respectively. Five species are consistently associated with each phenotype in all three countries. Specifically, *P. vulgatus, G. qucibialis, A. putredinis, Agathobacter rectalis,* and *Faecalibacterium prausnitzii D* are associated with strain coexistence, and *P. vulgatus, P. distasonis, Prevotella copri B, Bacteroides xylanisolvens,* and an unclassified Phocaeicola species are associated with species abundance (FigureS6, Extended Data2). In the Israeli cohort, participants carrying the *P. vulgatus* “blue” sub-type have 0.32% more variable positions and 0.0041 higher abundance than those with the “green” sub-type, with similar patterns observed in the other countries (Figure5).

Next, we leveraged the Israeli longitudinal data to test whether *P. vulgatus* strain turnover is associated with the host and bacterial characteristics. We find that individuals who experienced strain turnover in this species as they got four years older had a smaller increase in BMI (q<0.01) and a greater increase in species abundance (q<1e-4) compared to those in whom the strain persisted, independent of their baseline sub-type. Similar trends were seen after two years.

In conclusion, bacterial intra-species genetic variations capture fine-scale bacterial and host associations that may be missed at the species level, highlighting the importance of strain-level resolution in microbiome research.

### Bacterial subspecies associate with host physiology

To extend the analysis of subspecies associations beyond host age, sex, and BMI, we leveraged the extensive phenotypic data of the Israeli cohort^12^, comprising thousands of clinical, molecular, and behavioral host features previously grouped into 14 physiological systems^32^. For instance, the cardiovascular system was assessed based on blood pressure from monitors, blood pressure ratios computed in the ankle–brachial index (ABI) test, arterial stiffness estimated by pulse wave velocity (PWV), carotid intima–media thickness (IMT) computed from carotid ultrasound, retinal vascular parameters computed from retinal imaging, and electric activity of the heart as captured by electrocardiogram (ECG). For each of 96 species with identified sub-types across these physiological systems, we trained a gradient boosting decision tree model to predict the bacterial sub-type from host features (1,317 tested associations, methods). We identified 88 associations where model performance exceeded that of models trained on permuted bacterial sub-types, as well as baseline models using only host age and sex as features (Figure6, Extended Data3). Notably, 31 (35%) of the associations remain significant after adjusting for species abundance, indicating that sub-type effects are not only driven by varying species quantity. As expected, diet and serum metabolites were the most associated physiological domains, whereas Lachnospira were the most associated species. Interestingly, a previous study found unusually low SNP and gene coupling in these species^21^.

**Figure 6.**
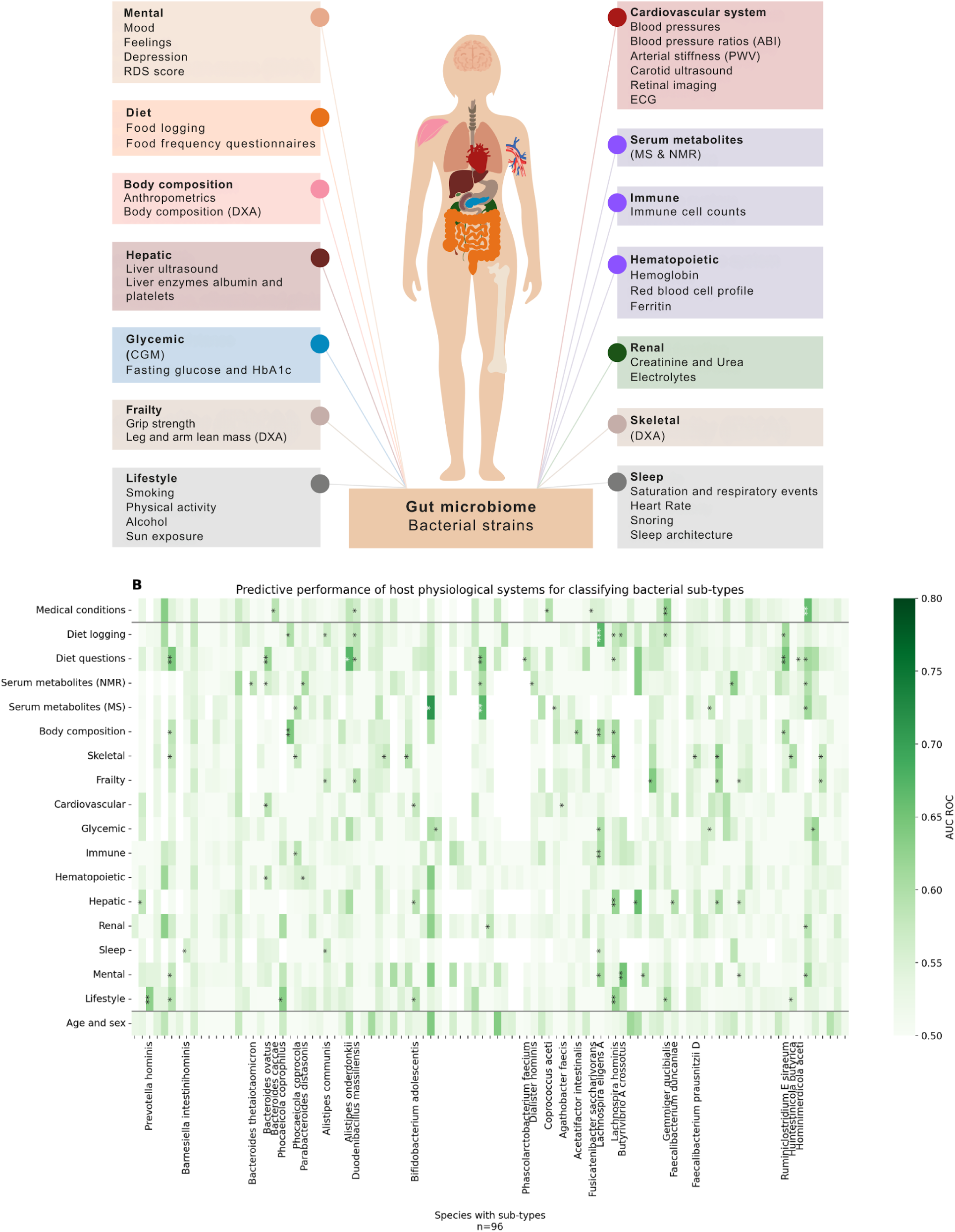
Bacterial subspecies associate with host physiology. **A.** Overview of the comprehensive clinical, molecular, and behavioral host data grouped into physiological systems. Adapted with permission from Reicher et al^32^. **B.** Predictive performance of host physiological systems (rows) for classifying sub-types of 96 bacterial species (columns). Performance is measured by the area under the receiver operating characteristic curve (AUC ROC) of a gradient boosting decision tree classifier. The top row represents medical conditions. Significance annotations indicate improvements in AUC ROC relative to models with permuted bacterial sub-types and baseline models using only host age and sex (bottom row): *>0.05, **>0.10, ***>0.15.

For example, sub-types of *Lachnospira eligens* (formerly *Eubacterium eligens*) showed associations with diet, glycemic control, body composition, immune function, sleep, and mental health. Key predictive features include intake of pizza, rice, and potatoes; estimated HbA1c; limb fat and bone mass; white blood cell count; sleep apnea indicators; and levels of professional, financial, and emotional satisfaction. Sub-types of *Lachnospira hominis* (also known as *Roseburia hominis*) were associated with diet, lifestyle, body composition, skeletal and hepatic health. Top predictive features include consumption of fruits, vegetables, and nuts; physical activity; femur and spine mass; liver attenuation; and serum bilirubin and albumin levels.

Importantly, these intra-species associations with host physiology were not limited to bacteria with specific subspecies population structures. For instance, *G. qucibialis*, which exhibits low LD, was linked to diet by beef and egg intake, lifestyle factors especially physical activity, serum metabolites, and medical conditions such as hemorrhoids and hyperlipidemia. *B. intestinihominis*, having moderate LD, was associated with sleep traits like oxygen desaturation. *P. distasonis*, which shows high LD, was tied to serum metabolites including alpha-lipoic acid (ALA), multiple high-density lipoprotein (HDL) particles, and hematopoietic markers mainly ferritin, hemoglobin, and red blood cells distribution width (RDW).

For a subset of 56 species and 12 physiological systems, we had sufficient follow-up data after two years to evaluate whether baseline bacterial sub-types associate with change in host physiological features (500 tested associations). Out of these, we identified 48 significant associations (Extended Data3), of which 21 (44%) remained significant after adjusting for species abundance. Interestingly, only three associations overlap with those previously identified using baseline physiological values. For example, *G. qucibialis* showed new associations with frailty (e.g., right-hand grip strength). *B. intestinihominis* became linked to lifestyle (physical activity and working hours), as well as serum metabolites and hepatic health (liver viscosity, attenuation, and elasticity). Similarly, *P. distasonis* was newly associated with changes in body composition (limb fat mass) and immune function (white blood cell count).

For an even smaller subset of 37 species, we had sufficient follow-up data after two years to evaluate whether bacterial strain turnover is associated with change in host physiological features (359 tested associations). Out of these, we identified 40 significant associations (Extended Data3), of which 26 (65%) remained significant after adjusting for species abundance. Interestingly, only four associations overlap with those previously identified using baseline bacterial sub-types. Notably, the hematopoietic system showed association with eight additional species. *Caccoplasma intestinavium,* which is not associated with any physiological system using baseline sub-types, is linked to five systems using strain turnover: diet, lifestyle, hematopoietic, immune, and renal function. In each of these systems, age is among the top predictive features. Nonetheless, system-specific features significantly improve prediction beyond a baseline model including only age and sex. Similarly, strain turnover in *L. hominis* introduced four new associations–with mental health, hematopoietic, immune, and renal function. In all of these cases, sex is the top predictor, yet system-specific features add significant value. Finally, strain turnover in *L. eligens* is linked to changes in cardiovascular features, consistent with its baseline links with diet, glycemic control, and body composition–all key determinants of cardiovascular health.

In conclusion, bacterial intra-species genetic variations carry rich phenotypic information and reveal specific associations with a wide range of host physiological domains, highlighting even more the relevance of strain-level resolution for understanding host–microbiota interactions.

## Discussion

Here, we analyzed the intra-species genetic diversity of 936 gut bacterial species across 24,997 individuals from Israel, the Netherlands, and the US. Across these three Western populations, bacterial parameters were remarkably consistent, suggesting that, within such settings, host genetics and geographic dispersal exert only a limited influence on bacterial population dispersal.

Strain-level analyses provided insights into bacterial population dynamics that extend beyond traditional such as species-level abundance. Whereas species-level measures showed reduced consistency across populations, kinships, and time–likely reflecting increased sensitivity to sampling time and sequencing protocols–whole-genome strain typing yielded more robust and informative patterns. Some of our findings have been previously reported (e.g.,^16,19–21,23,25,26^). However, the depth and breadth of our cohort^12^, including diverse human kinships within the same country, lifestyle, and age group, coupled with deep phenotypic and high-resolution longitudinal data spanning several years, allowed to refine, generalize, reduce bias, and strengthen previous findings for more species, in their natural habitat, than ever before.

For instance, gut strain richness has previously been characterized in cultivated species^23^ By systematically quantifying multi-strain colonization in vivo, we find clonal strains are often mutually exclusive, while genetically variable strains tend to coexist within the same host and be more prevalent across hosts. Species with strain coexistence tendencies carry quorum sensing, secretion systems, and a variety of metabolic genes in reference genomes, supporting a model in which inter-bacterial communication, genetic exchange, and metabolic niche partitioning enable stable multi-strain colonization within and across hosts.

From the bacterial perspective, highly abundant species such (e.g., Bacteroidota and Proteobacteria) exhibit greater strain stability, whereas low-abundance species (e.g., Firmicutes and Actinobacteria) display increased in-clonal mutations. Importantly, strain turnover within individuals over time is linked to the presence of chemotaxis and flagellar genes in reference genomes, as well as to species prevalence across individuals. This suggests a role for bacterial motility in strain acquisition from the host environment. Valles-Colomer et al.^26^ reported an opposite association with mobility in mother–infant pairs using a marker-gene approach. A possible explanation is that chemotaxis, which can help bacteria evade and modulate host immunity and is often associated with pathogenicity^33^, facilitates strain transmission in the general population, whereas maternal and infant mechanisms specifically restrict transmission of highly motile, potentially pathogenic, strains.

From the host perspective, gut bacterial strains were largely stable over several years, with only gradual and modest changes. Environmental elements, such as cohabitation, shared lifestyle, and bacterial richness, showed a stronger impact on strain sharing than host genetics. Notably, bacterial stress-response mechanisms, especially sporulation, were negatively associated with strain sharing among cohabiting partners. In contrast, Valles-Colomer et al.^26^ reported a positive association between sporulation and strain sharing among non-cohabiting individuals in the same population. This suggests long but not short-range strain transmission requires powerful survival mechanisms such as sporulation.

Bacterial intra-species genetic variations are associated not only with basic host characteristics such as age, sex, and BMI, but also with diverse physiological domains, like frailty. To assess the robustness of these findings, we compared our main results with existing literature. *P. vulgatus* stood out as it was consistently linked to all three host characteristics across countries and also to bacterial characteristics such as strain coexistence and species abundance. At the species level, *P. vulgatus* has been previously associated with obesity in both positive^34–37^ and negative^38–42^ directions. At the subspecies level, Zahavi et al.^6^ reported 909 SNPs in *P. vulgatus* associated with BMI, most in LD, with the top SNP in *arnC* corresponding to a 1.3 kg/m² BMI difference. *arnC* encodes a glycosyltransferase that modifies arabinose before its attachment to lipid A, the most immunogenic component of lipopolysaccharide (LPS)^43^. LPS is a well-established immunogenic driver of obesity phenotypes in mice^44^ and also serves as a receptor for many bacteriophages^45^. Such LPS variation can modulate both host inflammatory responses and phage–bacteria dynamics. Our study shows *P. vulgatus* strain turnover is associated with a smaller increase in BMI and a larger increase in species abundance compared to strain persistence. Taken together, our observations align well with the literature and suggest that *P. vulgatus* intra-species genetic variations and their turnover, rather than species abundance alone, may underlie conflicting obesity associations previously reported.

Lachnospira provides another illustration of how subspecies resolution sharpens host–microbiota links. *L. eligens* was associated with seven physiological domains and *L. hominis* with nine. Both species related to diet, body composition, immune function, and mental health; in addition, *L. eligens* associated with glycemic status, cardiovascular health, and sleep, whereas *L. hominis* associated with lifestyle, skeletal, hematopoietic, renal, and hepatic health. These patterns align well with prior work. Pascale et al.^46^ describes L. eligens is selectively stimulated by pectin, a plant cell-wall polysaccharides, particularly from potato fiber, using in vitro fermentation. In line with this, potato and pizza (that contains tomato pectin) intake were among the strongest dietary predictors of L. eligens sub-types in our models. Other studies linked L. eligens with lower postprandial glucose and insulin, reduced body weight and waist circumference, lower cardiovascular risk, and anti-inflammatory effects^46^. Vogl et al.^47^ utilized phage immunoprecipitation assay and found altered antibody responses to Lachnospiraceae flagellins and a protein involved in cell wall modifications in severe chronic fatigue syndrome. No species-level dysbiosis was found, consistent with strain-specific immune recognition. Conversely, L. eligens abundance has been positively associated with insomnia propensity in adults, yet negatively associated with childhood obstructive sleep apnea and adult depression^48–50^. Given the frequent co-occurrence of sleep impairment and depression, such mixed results are difficult to interpret at the species level. Our findings suggest that intra-species genetic variations in Lachnospira can help reconcile conflicting associations at the species level and direct functional experiments.

*P. distasonis* illustrates how our findings generate new testable mechanistic hypotheses. At the species level *P. distasonis* has been implicated in both pro- and anti-inflammatory across diseases, including obesity, where iron metabolism was shown to play a role^51^. In our study, *P. distasonis* sub-types were linked to change in host immune function and body composition after two years and were strongly predicted by baseline serum markers such as ALA, ferritin, hemoglobin, and RDW–all related to iron homeostasis^52^. These results suggest that specific P. distasonis sub-types may modulate, or are shaped by, host iron homeostasis, which in turn contributes to obesity, illustrating a novel testable mechanistic link emerging from our atlas.

Overall, our strain analyses provide insights into bacterial population dynamics that are not captured by the species level, and elucidate ecological and evolutionary principles governing strain stability, turnover, and coexistence in the human gut across more species than ever before. In parallel, our atlas linking strain-level genetic variation to 14 diverse host physiological systems identifies concrete targets for interventions such as diet, lifestyle, probiotics, and antibiotics. Together, the ecological and evolutionary principles (to predict and steer community responses) and the atlas (to select or engineer strains) could advance us towards personalized, microbiome-based interventions to improve human health.

### Study limitations

First, the data used in this study is derived from Western populations, which may limit the generalizability of the findings to non-Western lifestyles and environments. Addressing this will require similarly large, deeply phenotyped cohorts in underrepresented populations. To partially bridge this gap, the deep phenotypic cohort initiated in Israel is currently being expanded to other regions, including the United Arab Emirates and Japan^12^. Second, our analyses are influenced by variation in sample size across species, which may cause taxa such as *P. vulgatus* to appear frequently in our results. However, this reflects natural variation in prevalence and abundance in the human gut rather than a selection bias. Finally, the observational nature of our study precludes causal inference, preventing us from determining the directionality of the identified associations.

## Supplementary figures and table

**Table S1.** Accuracy of individual identification over time using different microbiome profiling methods. Percentage of correctly identified Israeli and Dutch individuals at two- and four-year follow-ups based on different microbial resolutions (strain or species) and feature types (existence or abundance).

|  |  |  |  |  |  |
| --- | --- | --- | --- | --- | --- |
| Follow-up [years] | 2 | 2 | 4 | 4 | 4 |
| Country | Israel | Israel | Israel | Israel | Netherlands |
| Method/<br>Metric [%] | Marker genes | Whole genome | Marker genes | Whole genome | Whole genome |
| Strain<br>existence | 84.63 | <u>98.07</u> | 68.72 | <u>94.93</u> | 92.77 |
| Strain<br>abundance | 46.46 | - | 28.05 | - | - |
| Species<br>existence | 84.00 | 86.59 | 67.21 | 71.74 | 75.30 |
| Species<br>abundance | 44.18 | 46.90 | 26.65 | 29.23 | 38.55 |

**Figure S1.**
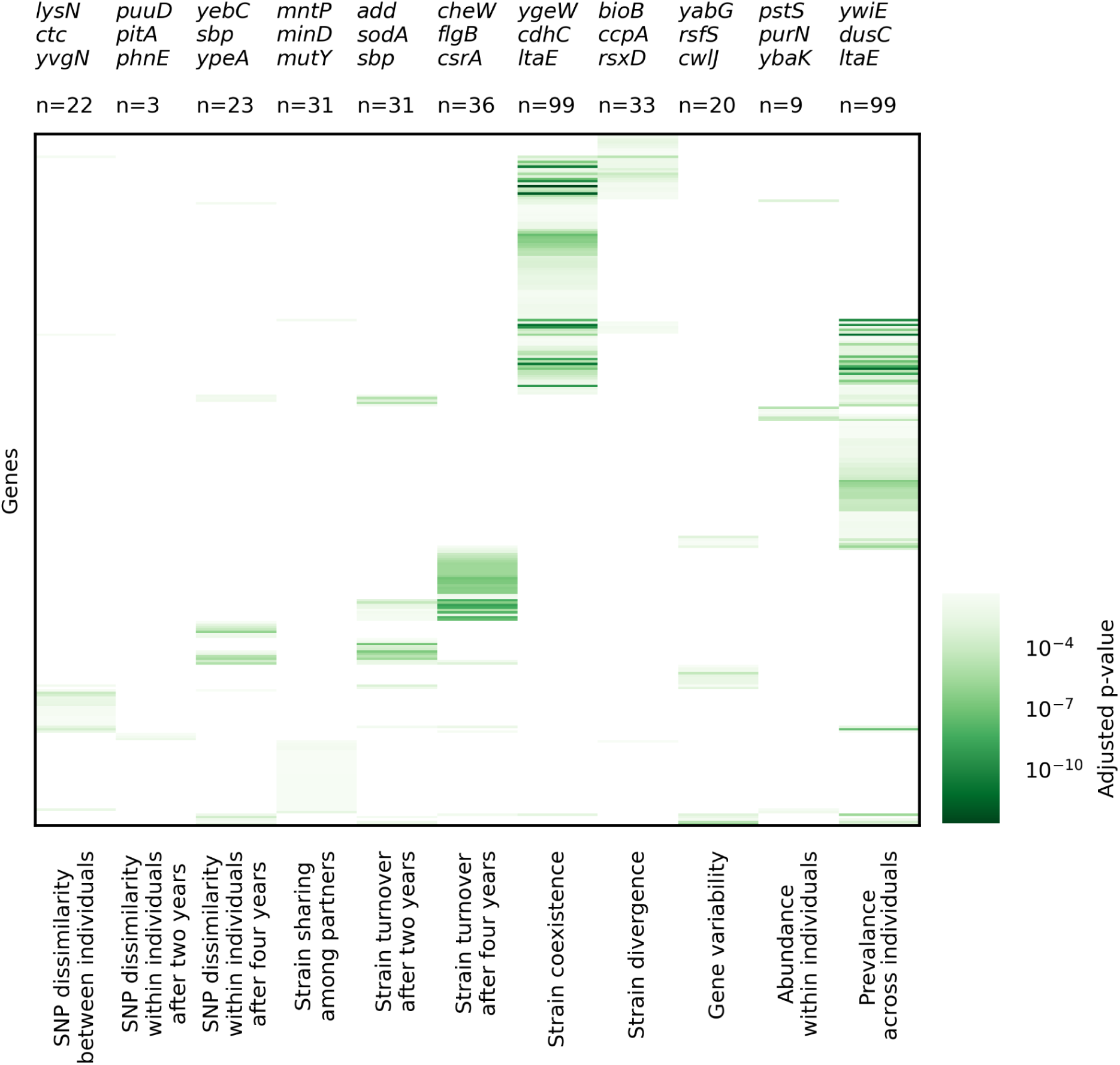
Functions associated with baseline characteristics of bacterial species. Cluster map of genes associated with bacterial parameters in the Israeli dataset (t test adjusted p<0.05, colorbar). The number of associated genes (n) and the top three genes per parameter are indicated above the plot. Abbreviations: SNP - single nucleotide polymorphism.

**Figure S2.**
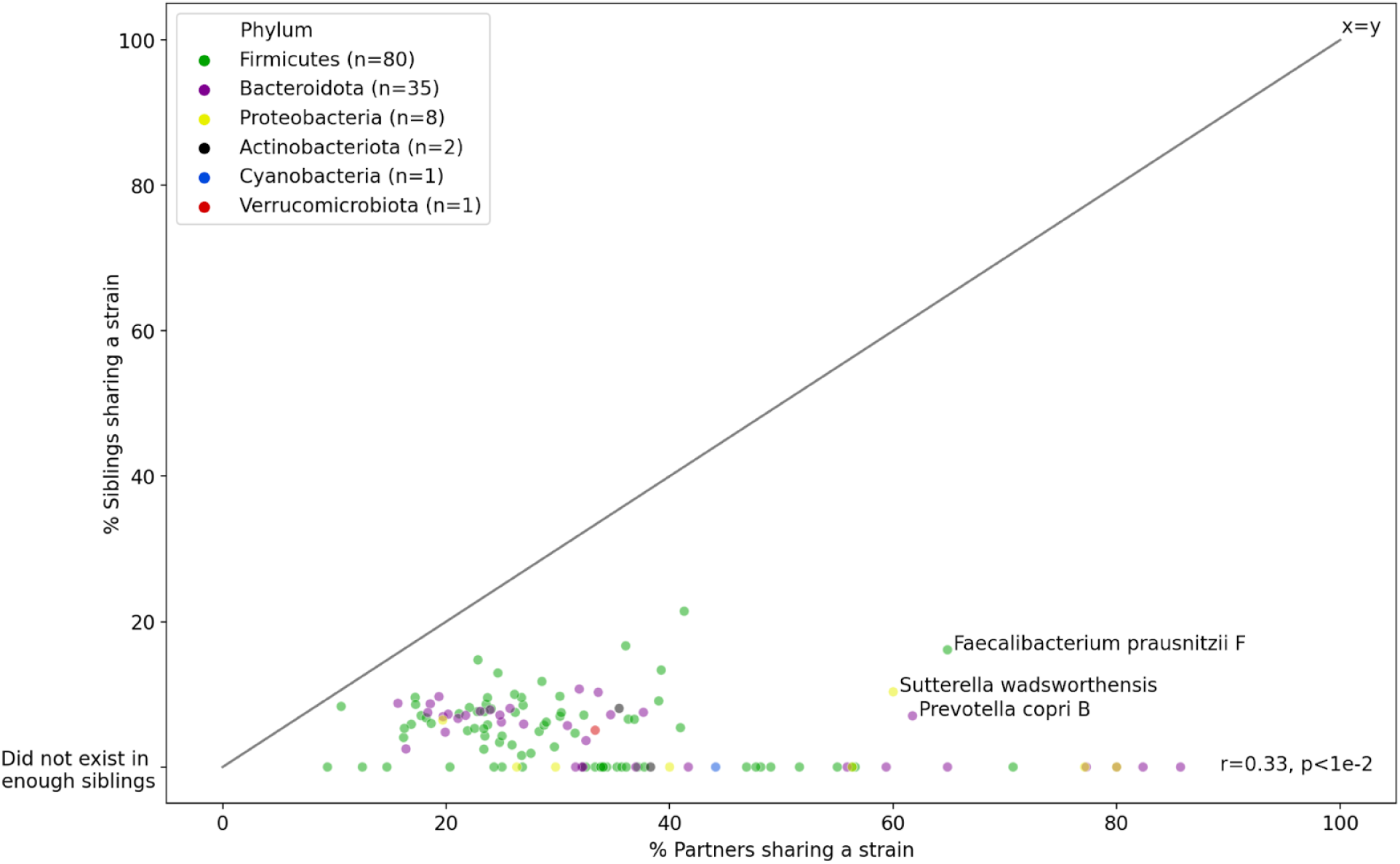
Environment dominates over host genetics in shaping human gut bacterial strains. Each dot represents a bacterial species, colored by its phylum (legend, n indicates number of species). The x-axis shows the percentage of cohabiting partners sharing a strain in each species, while the y-axis shows siblings.

**Figure S3.**
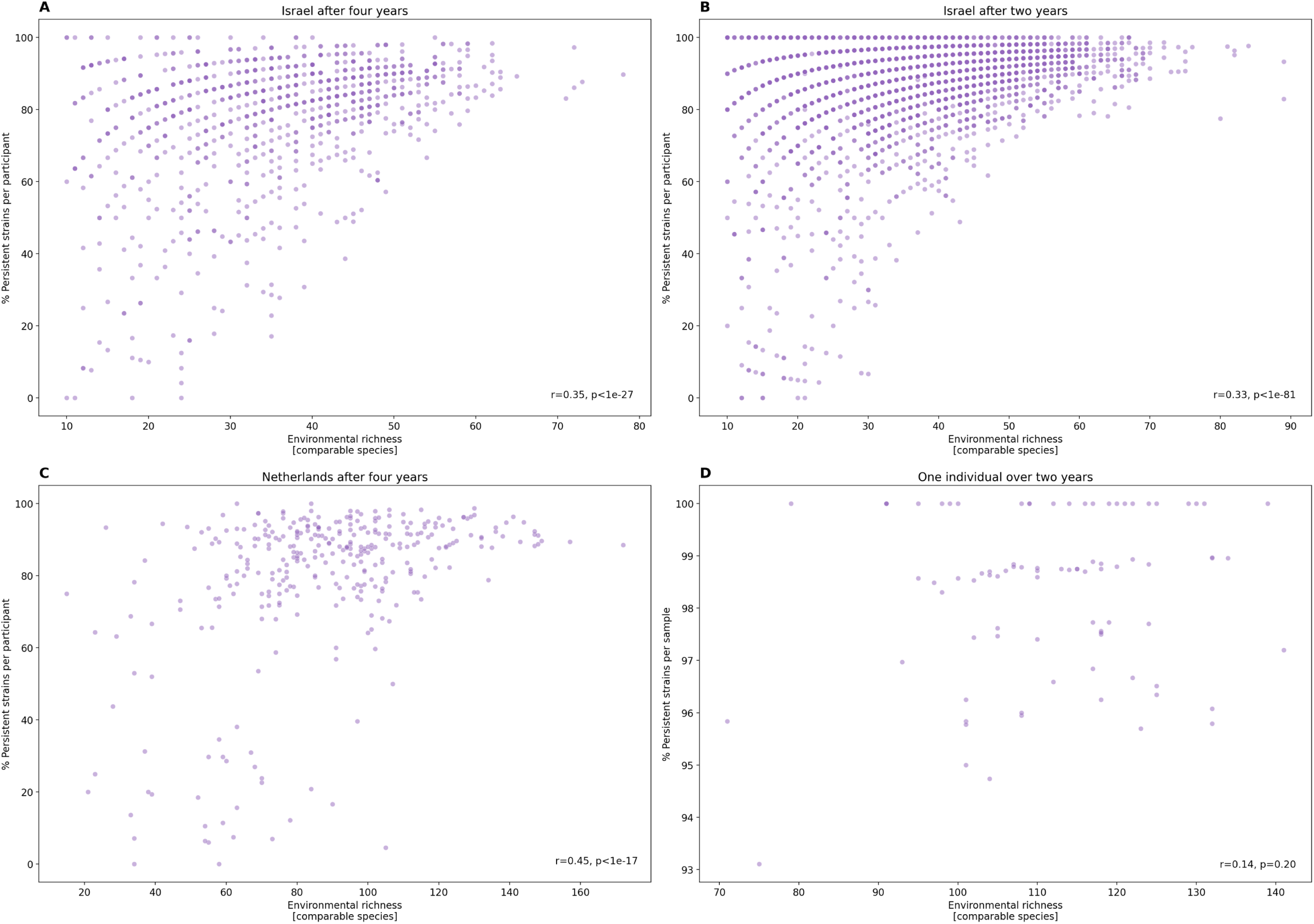
Individuals with greater bacterial richness exhibit higher strain stability. Each plot shows the relationship between the percentage of persistent strains (y-axis) and environmental richness portrayed by the number of comparable species (x-axis) for individual participants (dots). **A and B.** Israeli dataset after four and two years, respectively. **C.** Dutch dataset after four years. **D.** One Israeli individual over two years, with persistence measured relative to the previous sample. Pearson correlations are shown.

**Figure S4.**
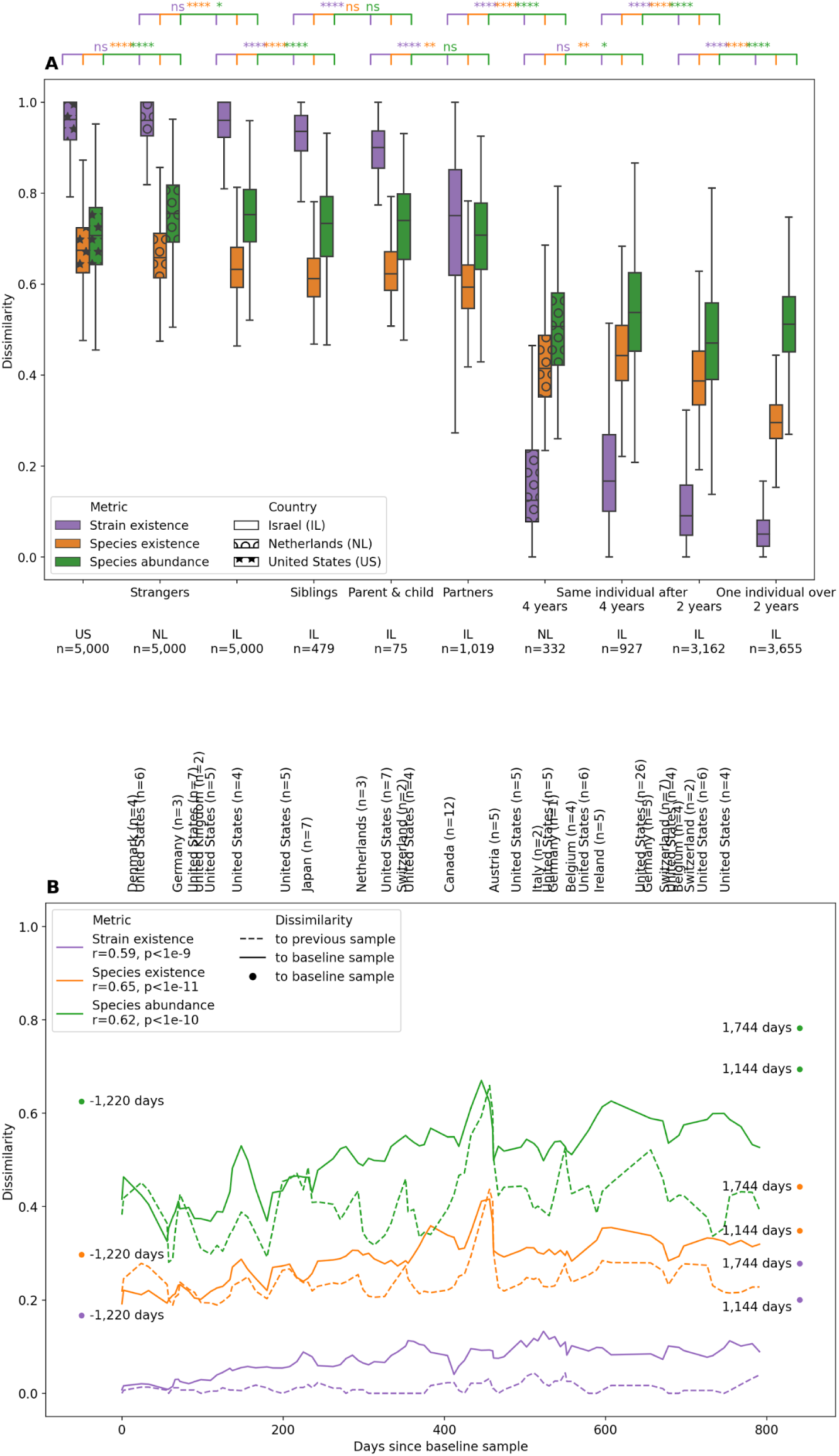
Strain typing is more robust than species composition. **A.** Strain-level dissimilarity (purple) is compared to species presence-absence (orange) and abundance (green) dissimilarities (y-axis) between individuals. Strangers are shown across three countries–Israel (IL, plain background), the Netherlands (NL, marked with circles), and the United States (US, marked with stars). Family members include siblings, parent–child pairs, and cohabiting partners. Within-individual comparisons are shown after four and two years, with the rightmost box representing one individual’s comparison of 86 samples. Boxes indicate quartiles; whiskers extend to 1.5 times the interquartile range. t tests q-value annotations: not significant (ns) q>0.05, *q<0.05, **<0.01, ***<0.001, ****<0.0001. **B.** Longitudinal data from a single individual across time (x-axis), showing a three-point rolling mean of the dissimilarity values (y-axis). Comparisons to the previous sample are shown as dashed lines; comparisons to the baseline sample are shown as solid lines. Three additional samples taken outside the two-year window are plotted as dots with their actual time relative to the baseline indicated. Pearson correlations between dissimilarities and days since the baseline sample are shown in the legend. International travels are marked above the plot, with trips duration (in days) indicated in parentheses.

**Figure S5.**
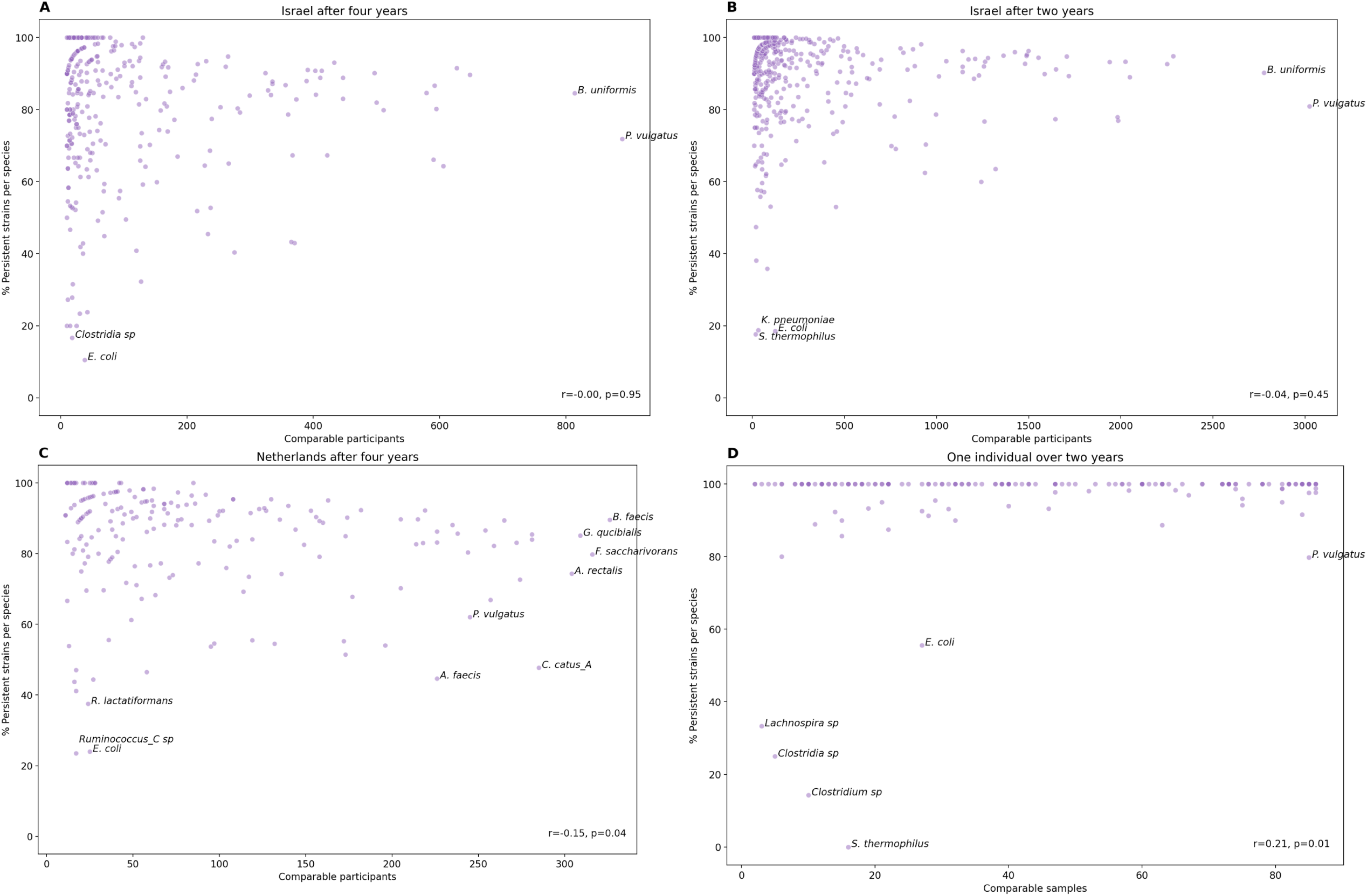
*Escherichia coli* and *Phocaeicola vulgatus* strains are highly dynamic. Each plot shows the relationship between the percentage of persistent strains (y-axis) and the number of comparable participants (x-axis) for each species (dots). **A and B.** Israeli dataset after four and two years, respectively. **C.** Dutch dataset after four years. **D.** One Israeli individual over two years, with persistence measured relative to the previous sample. Pearson correlations and labels of extreme species are shown.

**Figure S6.**
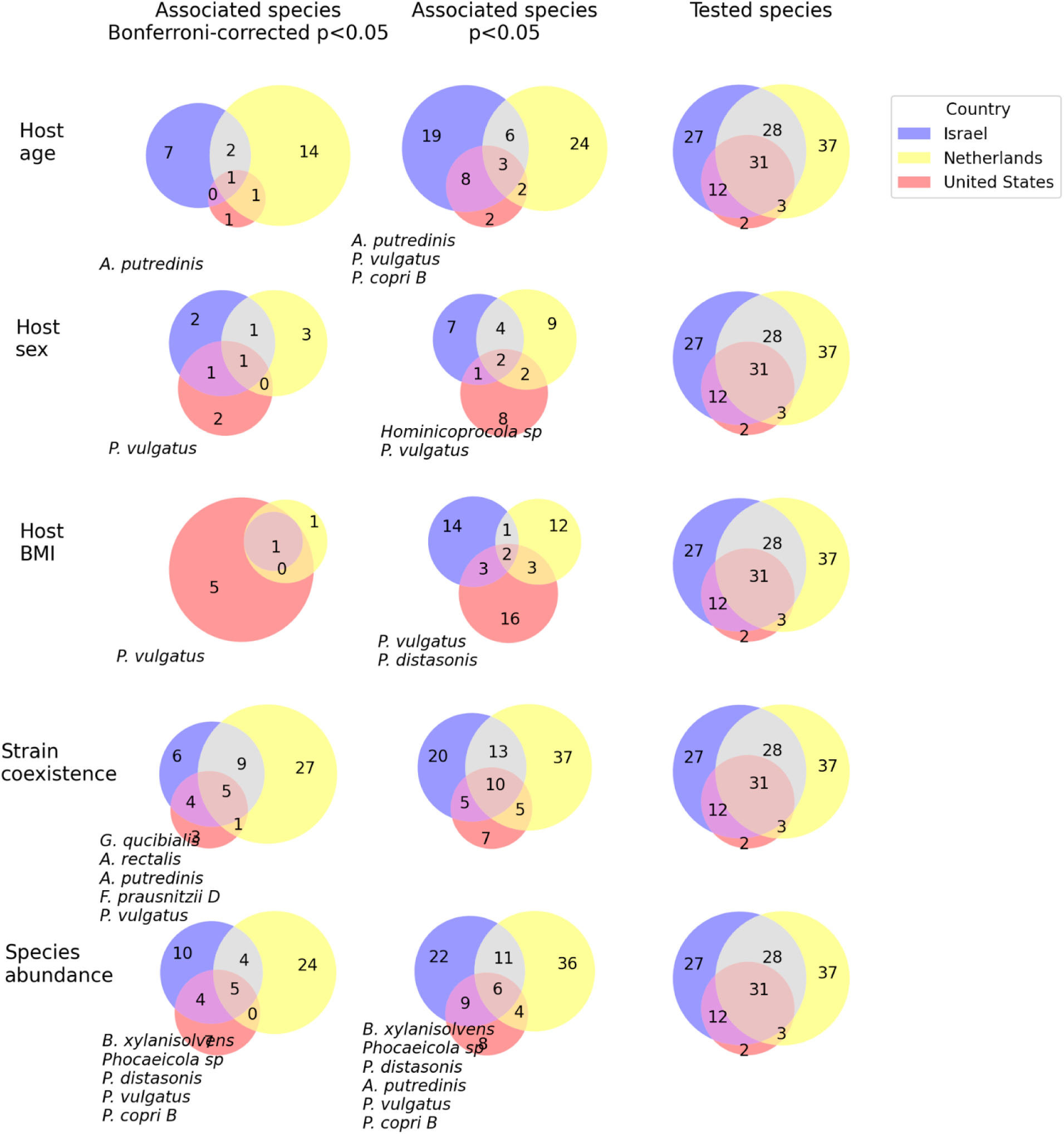
Bacterial subspecies associate with host age, sex, and BMI. Venn diagrams show the number of species tested, associated (linear models p<0.05), and significantly associated (q<0.05; columns) with host characteristics (age, sex, and body mass index–BMI) and bacterial parameters (variable positions indicative of strain coexistence and species abundance; rows), in the Israeli (blue), Dutch (yellow), and American (red) datasets (legend). If less than ten species replicate in all three countries, names are indicated.

## Methods

### Datasets

The main dataset used in this study is the Israeli Human Phenotype Project from Israel^12^, as of December 12, 2024. It included 9,600 stool samples from different individuals at their baseline visit, after excluding samples with <6M aligned reads, samples missing host age or sex, or with <50 detected species. A total of 706 species, each present in ≥500 individuals, were analyzed. An additional 3,220 two-year follow-up samples were available (2.14±0.22 years apart, range 1.03–2.99), as well as 948 four-year follow-up samples (4.10±0.22, range 3.07–4.94). Moreover, one individual contributed 86 stool samples over two years (9.24±6.72 days apart, range 1–49), and three additional samples outside this time period. In this individual, 339 species present in ≥20 samples were analyzed.

Two validation datasets were analyzed using the same filtering criteria as in the main dataset. The first validation dataset, Lifelines, was from the Netherlands. Lifelines-DMP^13^ included 8,178 stool samples from different individuals and 572 analyzed species. Lifelines-DEEP^14–16^ included 1,130 baseline samples from different individuals and 333 four-year follow-up samples (3.53±0.12 years apart, range 3.33–3.92). In this smaller dataset, 581 species present in ≥100 individuals were analyzed. The second validation dataset was DayTwo from the US^17^. It included 6,088 stool samples from different individuals and 505 species, meeting a filtering threshold of ≥4M aligned reads. The difference in filtering threshold for aligned reads stem from variations in read length and sequencing depth.

### Read processing

Metagenomic sequencing reads were processed using Trimmomatic to remove adapters and filter low-quality sequences (parameters: -phred33 ILLUMINACLIP::2:30:10 SLIDINGWINDOW:6:20 CROP:100 MINLEN:90)^53^. Human DNA was removed by aligning reads to the human genome (hg38) using Bowtie2^5^4. Remaining reads were aligned to the Weizmann Institute of Science (WIS) human gut microbial genome reference set (parameters: --score-min L,-40,0)^18^, with updated taxonomy according to the Genome Taxonomy Database (GTDB) version RS220^55,56^. Aligned microbial reads were retained for downstream analysis only if their highest-quality alignment score was unique to a single genomic location (parameters: min_base_quality=30). Finally, per-position coverage and allelic information were extracted using Bowtie2 “pile-up” command.

### SNP dissimilarity

Pairwise genetic dissimilarity between samples per species was quantified using Jaccard dissimilarity, defined as the proportion of genomic positions that lacked common alleles relative to the total number of comparable positions. Samples were compared only if ≥20K overlapping positions, each covered by ≥3 reads, were available. Low values were clipped to 1/20K, reflecting the method’s detection threshold. When computing the percentage of turnover, persistent, or shared strains between samples ≥10 coexisting species were required.

### Linkage disequilibrium

LD was calculated as the Pearson correlation squared between major allele frequencies of SNPs within the same contig, given that ≥50 SNPs were available in the contig. SNPs were included in the analyses if they were detected in ≥500 individuals, with a non-major allele present in ≥5% and ≥50 individuals in each dataset. The major allele was defined universally across datasets as the most common allele in 100K samples available in the lab. To optimize memory usage, allele frequencies were normalized as integers on a 0–200 scale, and to further reduce computational burden, 40% of correlations were randomly sub-sampled when computing LD.

### Gene variability

Gene variability was quantified as the exponent in Heap’s law^57^ in the WIS human gut microbial pangenome reference set^5^. The exponent signifies whether gene discovery has reached a plateau (0, minimal gene discovery) or it continues to grow as new strains are introduced (1, maximal gene discovery) to the pangenome.

### Species composition

Species abundance was estimated using the Unique Relative Abundance (URA) technique (parameters: num_mapped_to_subsample=6M, min_mapped_to_retain=1M, min_partial=0.5)^17^, which calculates the normalized mean coverage over the 50% most densely covered regions of each species regenome. To account for detection limits, low values were clipped to 1/10K. When computing Bray-Curtis and Jaccard dissimilarities over species relative abundance and existence respectively, only samples containing ≥10 coexisting species were compared.

### Subspecies

Sub-types were identified within species with ≥500 baseline samples (and *E. coli*) using average Euclidean clustering of their SNP dissimilarity. During the clustering step, samples with ≤20% missing dissimilarity values were temporarily imputed using the median or otherwise excluded. Species sub-types were defined as primary groups from the apex of the species dendrogram, displaying genetic divergence of ≥1% from the parent branch and containing ≥10% but ≤80% samples.

### Association with bacterial functions

Associations between bacterial parameters and functions were assessed using annotated genes published with the WIS human gut genome reference set^18^, disregarding copy number variations, present and absent in ≥30 species.

### Association with host characteristics

The association between species sub-types and host age, sex, and BMI in the three countries was assessed using Ordinary Least Squares (OLS) and Logit models, respectively.

To analyze the broader phenotypic associations, data from the Israeli dataset was divided into physiological systems, as described in Rechier et al^32^. For the 96 species with 2-5 identified sub-types, an eXtreme Gradient Boosting (XGB) classifier was trained to predict the type based on host age, sex, and physiological system features. Data was split into 80% training and 20% testing, with a five-fold cross-validation scheme employed within the training set (parameters: eval_metric=mlogloss, scoring=roc_auc_ovr, n_estimators=1000, max_depth=2). During training, a hyperparameter grid search was conducted over a small set of parameters (parameters: learning_rate=[0.01, 0.05, 0.001], min_child_weight=[20, 30, 40], colsample_bytree=[0.1, 0.2], subsample=[0.4, 0.6, 0.8]). To ensure robustness, the training process was repeated five times with different random seeds, allowing confidence intervals to be computed for the performance on the same test set. All steps were stratified by the species sub-type to maintain consistent class proportions. Feature importance was estimated using SHapley Additive exPlanations (SHAP) in one of the trained models.

## Acknowledgements

We thank members of the Segal, Pilpel, and Pollard groups for fruitful discussions and Liron Zahavi and Veronika Dubinkina for their valuable feedback on the manuscript. We thank Lifelines and Groningen Microbiome Hub team for making their large metagenomics studies publicly available. S.S is supported by the Israeli Council for Higher Education (CHE) via the Weizmann Data Science Research Center. E.S. is supported by the Crown Human Genome Center, and grants funded by the Israel Science Foundation (ISF). This study was supported by the National Heart, Lung, and Blood Institute (NHLBI) grant #R01-HL160862.

## Author contributions

S.S. conceived, designed, and performed the analysis, analyzed the results, and wrote the paper. A.G. processed the data. A.W. developed protocols and oversaw sample collection and processing. K.P., Y.P., and E.S. supervised the study.

